# A comparative study between a power and a connectivity sEEG biomarker for seizure-onset zone identification in temporal lobe epilepsy

**DOI:** 10.1101/2023.11.23.568472

**Authors:** Manel Vila-Vidal, Ferran Craven-Bartle Corominas, Matthieu Gilson, Riccardo Zucca, Alessandro Principe, Rodrigo Rocamora, Gustavo Deco, Adrià Tauste Campo

## Abstract

**Background:** Ictal stereo-encephalography (sEEG) biomarkers for seizure onset zone (SOZ) localization can be classified depending on whether they target abnormalities in signal power or functional connectivity between signals, and they may depend on the frequency band and time window at which they are estimated.

**New method:** This work aimed to compare and optimize the performance of a power and a connectivity-based biomarkers to identify SOZ contacts from ictal sEEG recordings. To do so, we used a previously introduced power-based measure, the normalized mean activation (nMA), which quantifies the ictal average power activation. Similarly, we defined the normalized mean strength (nMS), to quantify the ictal mean functional connectivity of every contact with the rest. The optimal frequency bands and time windows were selected based on optimizing AUC and F2-score.

**Results:** The analysis was performed on a dataset of 67 seizures from 10 patients with pharmacoresistant temporal lobe epilepsy. Our results suggest that the power-based biomarker generally performs better for the detection of SOZ than the connectivity-based one. However, an equivalent performance level can be achieved when both biomarkers are independently optimized. Optimal performance was achieved in the beta and lower-gamma range for the power biomarker and in the higher-gamma range for connectivity, both using a 30 s period after seizure onset.

**Conclusions:** The results of this study highlight the importance of this optimization step over frequency and time windows when comparing different SOZ discrimination biomarkers. This information should be considered when training SOZ classifiers on retrospective patients’ data for clinical applications.

## 1 Introduction

Presurgical evaluation of drug-resistant epilepsy often involves the use of intracranial EEG recordings to accurately localize the seizure onset zone (SOZ) (Bancaud, 1980). Recordings from hundreds of sensors are continuously acquired for up to three weeks and visually inspected by highly specialized neurologists. This can be a very challenging procedure given that seizure onset patterns may involve several frequencies (Perucca et al., 2014; Lagarde et al., 2016; Vila-Vidal et al., 2020b) and may have complex spatial distributions (Bartolomei et al., 2017).

Several stereo-encephalography (sEEG) biomarkers have been developed to help localize the SOZ using ictal activity. These biomarkers can be classified into two groups, depending on their target features: power or connectivity (for an extensive review see Vila-Vidal and Tauste Campo (2023)). On one hand, power-based biomarkers target abnormal activity patterns that might correlate with the SOZ. These methods are based on a spectral analysis of the sEEG signals of each region, from which changes or activations at specific frequencies are then extracted and evaluated. Some methods within this class, such as the Epileptogenicity Index (Bartolomei et al., 2008a), rely on detecting changes at specific frequency bands, often in the beta and gamma ranges (David et al., 2011; Murphy et al., 2017). Alternative methods offer more flexibility and can adapt to or combine different frequencies of interest (Gnatkovsky et al., 2011, 2014; Vila-Vidal et al., 2017, 2020b).

On the other hand, connectivity-based biomarkers target abnormal changes in functional connectivity, i.e., statistical dependencies of signals between pairs of electrodes. The rationale behind this approach is based on a conceptualization of epilepsy as a disease affecting a network of interconnected regions that generate and propagate ictal activity across the brain (Bartolomei et al., 2017). Different types of statistical relationships (eg. linear, non-linear, phase-based, etc.) have been used to build biomarkers that capture alterations in these functional relationships during ictal periods (Gotman and Levtova, 1996; Bartolomei et al., 2004; Mierlo et al., 2013; Nahvi et al., 2023; Balatskaya et al., 2020).

The studies mentioned above built their biomarkers based on either power or connectivity alone. One of the few exceptions in this regard is the work done by (Balatskaya et al., 2020), which combined the two types of measures to define a joint biomarker. A major limitation of this study, however, was the use of a predefined frequency band (beta-gamma range) and a predefined period of interest (20-30 s after seizure onset), which may fall short of capturing relevant activity given the heterogeneity observed across seizures (Perucca et al., 2014; Lagarde et al., 2016). Up to now, no study has compared the performance of these two type of biomarkers as a function of signal parameters such as the frequency and the time windows of interest.

The present study aimed to systematically compare the performance of a power– and a connectivity-based biomarker to identify SOZ contacts from ictal sEEG recordings. As a general power-based measure, we used the normalized mean activation (Vila-Vidal et al., 2017), which quantifies the average power activation with respect to baseline activity in a frequency band and time window of interest and can be regarded as a time-frequency generalization of previous known biomarkers (Bartolomei et al., 2008b; Gnatkovsky et al., 2014). As a representative of connectivity-based biomarkers, we defined an analogous measure relying on the cross-correlation (Kramer et al., 2008; Mierlo et al., 2014; Khambhati et al., 2017). The normalized mean strength (nMS) corresponds to the mean functional connectivity (cross-correlation) of every contact with the rest of signals in a frequency band and time window of interest.

The optimal frequency bands and time windows for identifying the SOZ might vary between power and connectivity-based measures. To account for these potential differences, we first evaluated statistical discrimination using Cohen’s d across different frequency bands and time windows of interest. To conduct a detailed examination of the classification performance of each measure, four criteria were used to select parameter combinations that maximized the classifying potential of each measure individually. The initial two criteria relied on the area under the curve (AUC) of the receiver operating characteristic (ROC), ensuring effective classification performance. The remaining two criteria were based on the F2-score, which prioritizes recall over precision (Yuan et al., 2024; Semeux-Bernier et al., 2023) to provide binary predictions (SOZ/non-SOZ) per contact.

This method was tested with a dataset of 67 seizures from 10 patients diagnosed with pharmacoresistant temporal lobe epilepsy (TLE) that underwent sEEG evaluation during pre-surgical diagnosis (Talairach et al., 1974; Munari and Bancaud, 1985; Guenot et al., 2001; Cardinale et al., 2013). The application of this comparative study to a TLE dataset was mainly motivated by an own previous publication (Vila-Vidal et al., 2017), in which we demonstrated in a subsample of the present cohort of TLE patients that this class of patients exhibited individually a similar normalized power quantification across seizures. Consequently, this was instrumental to perform averages of quantifiers across seizures and define one biomarker per patient.

## 2 Methodology

### 2.1 Patients and recordings

In this study, we analyzed sEEG recordings from a total of 67 seizures from 10 patients with pharmacoresistant focal-onset epilepsy who underwent presurgical evaluation at the Epilepsy Centre of Hospital del Mar (Barcelona, Spain) between 2012 and 2017. Seizure onset and termination times were independently marked by two epileptologists (RR and AP) using standard clinical assessment. For each seizure we analyzed sEEG recordings from the marked ictal epoch together with 60 s of pre-ictal and 60 s of post-ictal epochs. In this study, we included only patients in which the seizure focus had been marked by epileptologists under the general principle that “the ictal onset was confined to a certain number of contacts and it was stable through ictal events”. Patients’ characteristics are summarised in Table 1.

**Table 1.**
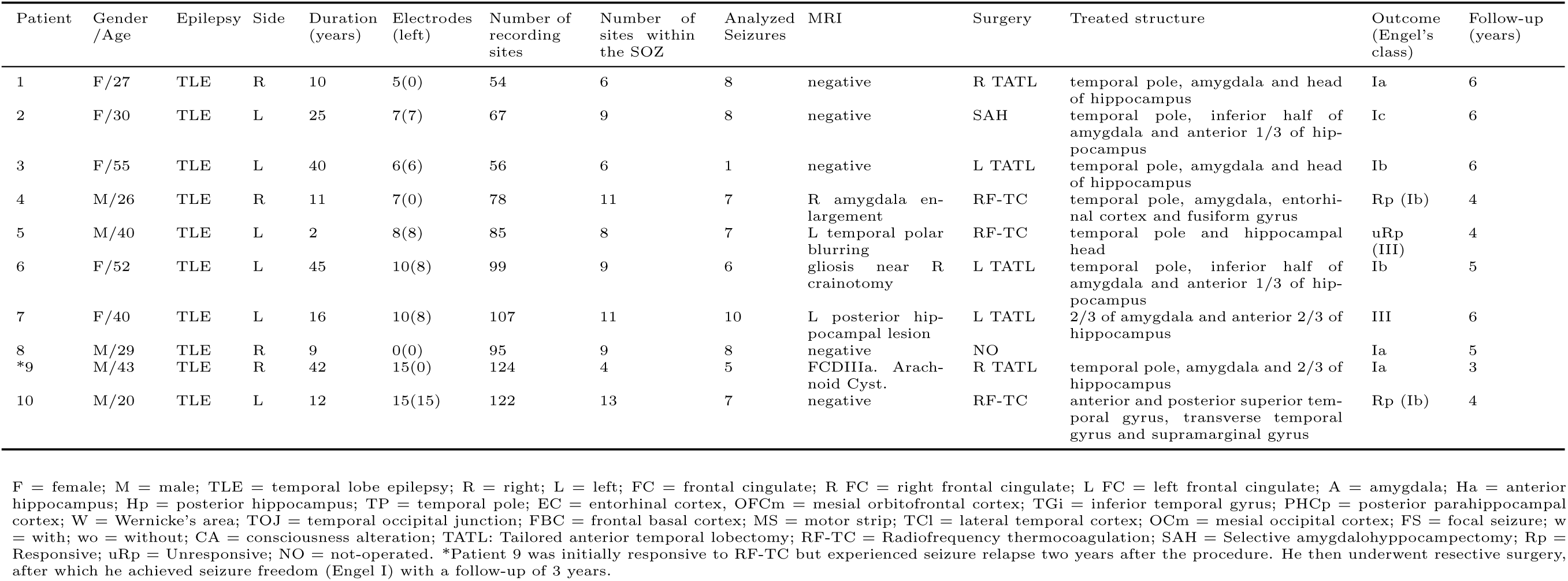
Main data of patients included in the study.

Stereo-EEG monitoring was performed using a standard clinical intracranial EEG system (XLTEK, a subsidiary of Natus Medical) with a sampling rate of 500 Hz,(in patient P3, the sampling rate was 250 Hz). Recordings were obtained using intracranial multichannel electrodes (Dixi Medical, Besaņcon, France; diameter: 0.8 mm; 5–15 contacts, 2 mm long, 1.5 mm apart) that were stereotactically inserted using robotic guidance (ROSA, Medtech Surgical, Inc). The decision to implant, the selection of the electrode targets and the implantation duration were entirely made on clinical grounds.

### 2.2 Data preprocessing

EEG signals were processed in the referential recording configuration (i.e., each signal was referred to a common reference). The electrodes per patient included in the analysis are reported in Table 1. We visually identified noisy electrodes and removed them from the analysis. Signals were band-pass filtered between 1 to 150 Hz using a zero-phase FIR filter (53 dB stopband attenuation, maximal ripples in passband 2%) to remove slow drifts and aliasing effects. A notch filter was applied to 50 Hz and its multiples to remove the power line interference (band-stop FIR filter with band-stop width of 1/10 and 1/35 of the central frequency for 50 Hz and its harmonics, respectively; 53 dB attenuation at center frequency, maximal ripples in passband 2 %).

### 2.3 General procedure

Our approach was to define and compare two biomarkers, one based on signal power and the other based on functional connectivity, to identify the seizure onset zone in the cohort of patients described above. The first biomarker used in this study is the previously introduced normalized mean activation (nMA) (Vila-Vidal et al., 2017, 2020b). The nMA quantifies the average power activation of each contact with respect to baseline activity within a frequency band and time window of interest. Similarly, we defined the normalized mean strength (nMS) to quantify each contact’s average functional connectivity strength (i.e., its average functional connectivity to all other contacts) over the same time-frequency ranges. The rationale is that, in parallel to putative changes in signal power at electrode contacts, the statistical dependencies of signals between pairs of contacts may change independently as a function of each contact’s location with respect to the SOZ (Bartolomei et al., 2004; Varotto et al., 2012; Mierlo et al., 2013). In addition, these changes are known to evolve over time (Courtens et al., 2016) and by using different temporal windows we may be able to capture different stages of the seizure start before propagation.

Both biomarkers are thus expected to depend on frequency bands and time windows and thus their respective level of SOZ discrimination might be optimized over the timefrequency space. We defined 6 frequency bands of interest (FOIs) and 9 time windows of interest (TOIs) to account for this parameter-dependence. FOIs were defined based on canonical bands: broadband (3-160 Hz), *δ*-*θ* (3-8 Hz), *α* (8-12 Hz), *β* (12-30 Hz), low-*γ* (30-70 Hz) and high-*γ* (70-160) Hz. All time windows of interest started at the annotated seizure onset time (reference time, 0 s) and were defined by 9 different window lengths: 0-1 s, 0-2 s, 0-3 s, 0-4 s, 0-5 s, 0-10 s, 0-20 s, 0-30 s and whole seizure. Seizure onset zone discrimination was independently evaluated for all possible FOI-TOI combinations with nMA and nMS. The processing steps for each biomarker are described in the following paragraphs.

### 2.4 Power-based biomarker: normalized mean activation

The processing used to compute the normalized mean activations (nMA) in each FOI and TOI was very similar to previous works (Vila-Vidal et al., 2017, 2020b). In brief, signals were first filtered in 42 non-overlapping logarithmically-spaced narrow frequency bands, each with a bandwidth of 10% of its lower frequency bound, covering the whole spectrum of interest 3-160 Hz. The Hilbert transform was then used to obtain the continuous power in each narrow band. Summation of signal power across narrow bands was used to obtain the total power of each contact in each FOI. Note that using a fixed bandwidth of 10% determines the cutting points of each band, which causes the resulting bands to have small deviations from the predefined FOIs. The resulting cutting points were: 3, 7.8, 12.5, 29.5, 69.7, 160 Hz. Following (Vila-Vidal et al., 2017), we used a preictal baseline of activity from 60 to 20 s before ictal onset. Baseline power values from all contacts were pooled together to build a baseline distribution. We then normalized each contact’s power with respect to this baseline and defined the instantaneous activation as the resulting z-score. This procedure was independently done for each FOI.

The mean activation (MA*_i,f,t_*) was then defined for each FOI *f*, TOI *t* and contact *i* as the average of its instantaneous activation in FOI *f* and across TOI *t*. This step was repeated for each seizure, thus obtaining a mean strength profile across contacts for each seizure, frequency, and time of interest. To allow for cross-seizure comparison, we normalized mean activation values within each seizure, frequency, and time of interest. We computed the mean *µ*_MA*,f,t*_ and standard deviation *σ*_MA*,f,t*_ of MA values in each frequency *f* and time window *t* across contacts. We then defined the normalized mean activation (nMA) for each contact *i*, FOI *f* and TOI *t* as the z-score the mean strength:

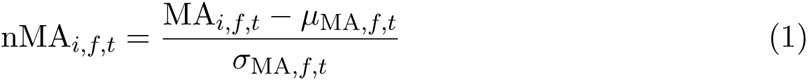

### 2.5 Functional connectivity-based biomarker: normalized mean strength

We first filtered each contact’s signal in the six FOIs using a zero-phase FIR filter (53 dB stopband attenuation, maximal ripples in passband 2%, Fig. 1A). We then estimated the functional connectivity between each pair of contacts in slicing windows of length 1s. Within each time window, the functional connectivity was estimated as the maximum of the absolute value of the cross-correlation for lags between –0.1 and 0.1s (Fig. 1B). Note that, besides its previous application as a functional connectivity estimator (Mierlo et al., 2014), the cross-correlation is a respresentative measure because its definition is analogous to a time-domain counterpart of the spectral coherence.

**Figure 1.**
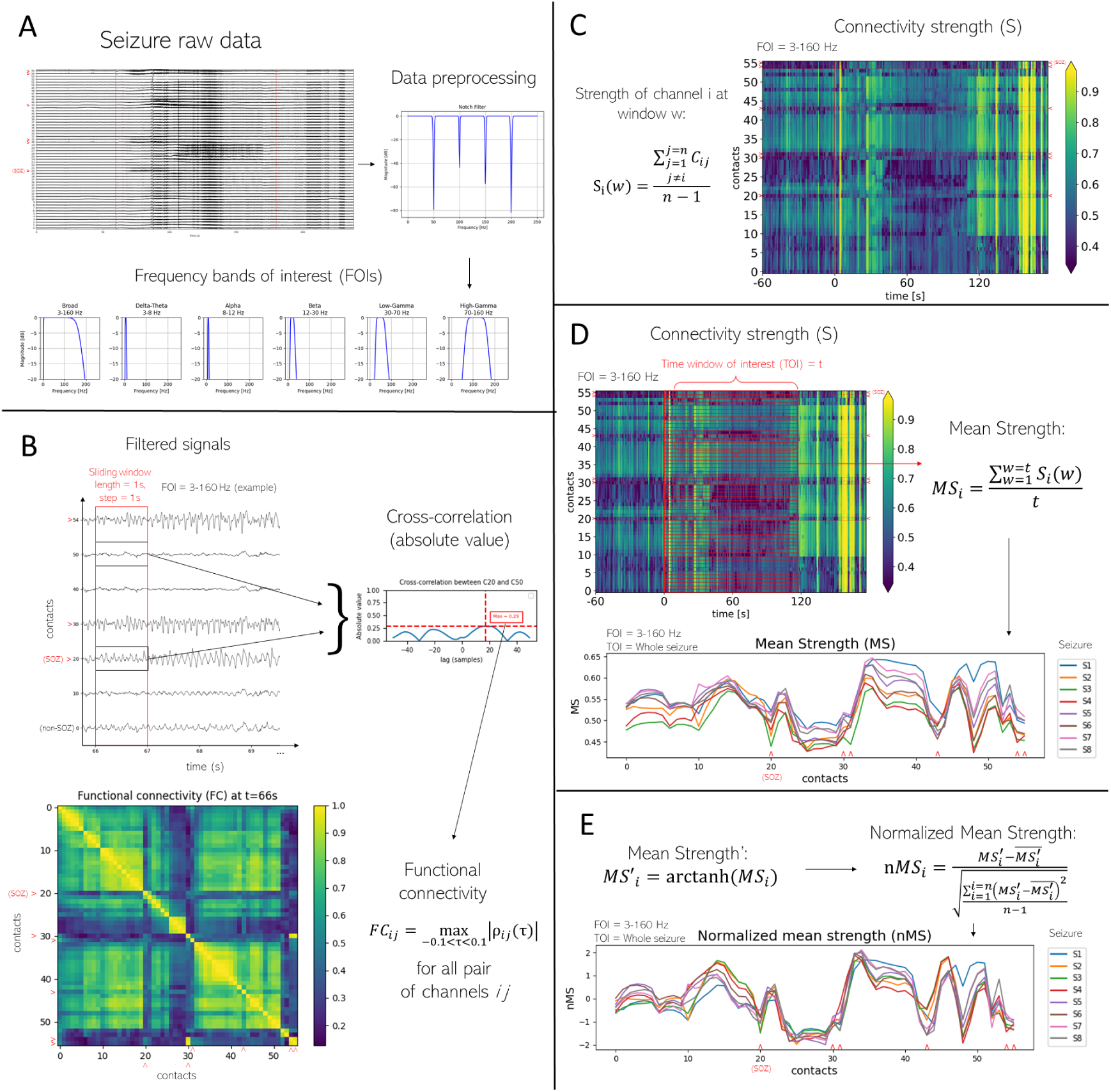
Functional connectivity-based biomarker: normalized mean strength (nMS) **A**. Data preprocessing and canonical band filtering. Raw signals were notch-filtered to remove the power line interference at 50 and its multiples. Vertical red lines mark seizure onset and offset. Seizure onset is used as the reference time (0 s). Signals were then band-pass filtered in six frequencies of interest (FOI), including 5 canonical bands and a broadband component: broadband: 3-160 Hz; *δ*-*θ*: 3-8 Hz; *α*: 8-12 Hz; *β*: 12-30 Hz; low-*γ*: 30-70 Hz; high-*γ*: 70-160 Hz. Artifacted noisy contacts were visually identified in this step and removed for the following steps. **B.** Time-varying functional connectivity. Using a sliding window approach (window length and step of 1 s), the time-varying functional connectivity (FC) was computed for each pair of contacts. The functional connectivity was estimated as the maximum absolute value of the cross-correlation across lags ranging from –0.1 to 0.1 s. **C.** Time-varying connectivity strength. The connectivity strength (S) was computed as the mean of the functional connectivity between each contact and the rest of the contacts for each time window. **D.** Mean strength in each time window of interest. For each contact and time window of interest (TOI), the mean connectivity strength (MS) was computed as the mean connectivity strength in the TOI. In particular, the following TOIs were explored: 0-1s, 0-2s, 0-3s, 0-4s, 0-5s, 0-10s, 0-20s, 0-30s, and whole seizure period. This procedure was repeated for each seizure. **E.** Normalized mean strength. To allow comparisons across seizures per patient, MS values were Fisher transformed and normalized (z-score) in each seizure separately, thus obtaining a normalized mean strength (nMS) for each contact over a given seizure. For each patient’s data, steps B-E were repeated for all FOIs and TOIs.

To demonstrate that there were neither volume conduction nor non-silent reference effects into the sEEG data, the maximum cross-correlation values along with their re-spective lags of all the SOZ/non-SOZ pair of contacts for windows spanning 5 seconds before and after seizure onset are shown in Fig. S1. The distribution of maximum values was observed to be predominantly below 1, and the lags were not uniformly 0, rejecting any substantial influence of volume conduction and other common reference effects into the obtained connectivity results.

We then defined a strength value (S) for each contact as the mean of its connectivity with all other contacts in each sliding window (Fig. 1C). The mean strength (MS*_i,f,t_*) was then defined for each FOI *f*, TOI *t* and contact *i* as the mean strength in frequency band *f* across time window *t*. This step was repeated for each seizure, thus obtaining a mean strength profile across contacts for each seizure, frequency and time of interest (Fig. 1C). To allow for cross-seizure comparison, we normalized mean strength values within each seizure, frequency, and time of interest. We first transformed all MS values using the Fisher z-transform (MS*^′^*) (Fisher, 1921). We then computed the mean *µ_f,t_* and standard deviation *σ_f,t_* of the transformed values in each frequency *f* and time window *t* across contacts. We then defined the normalized mean strength (nMS) for each contact, FOI *f* and TOI *t* as the z-score of its Fisher-transformed mean strength (Fig. 1E):

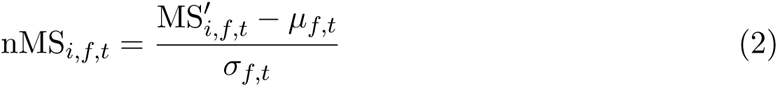

### 2.6 Statistical analyses

Before performing patient-level analysis, we aimed to assess homogeneity across seizures in each patient. Broadband nMA similarity across seizures was previously shown (Vila-Vidal et al., 2017). Here, we aimed to assess the similarity of nMA and nMS profiles across seizures within each FOI (using the whole seizure as the time window of interest). To do so, we computed the Pearson correlation of nMA between each pair of seizures for each FOI. We then average these values across all pairs of seizures to obtain a measure of inter-seizure similarity for each patient and FOI. The same procedure was used to quantify the inter-seizure nMS similarity. Next, we assessed the statistical power of these variables (nMA and nMS) to differentiate between SOZ and non-SOZ contacts at the patient level. To do so, we computed the median nMA (resp. nMS) across seizures and obtained a single value for each contact in each FOI-TOI combination. We then assessed the effect size of differences between SOZ and non-SOZ contacts using Cohen’s *d* (Cohen, 2016), which is defined as the difference between two means (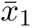 and 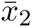) divided by the pooled standard deviation calculated from the variances (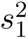 and 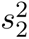):

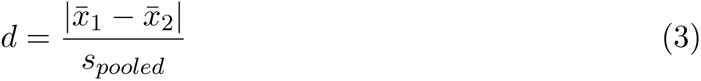

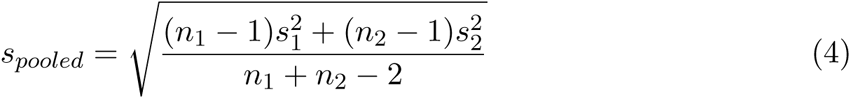

In addition, we also tested for statistical differences using a non-parametric test (Wilcoxon rank-sum test) and controlled the family-wise error rate in multiple comparsions using the Holm-Bonferroni step down procedure (Holm, 1979).

Finally, we compared nMA and nMS as features of a binary classifier for SOZ identification. Specifically, for each variable (nMA and nMS) we computed a receiver operating characteristic (ROC) curve and extracted the area under the curve (AUC) for each FOI-TOI combination. This procedure was done in each patient separately. We then aimed to maximize each classifier’s performance under two criteria. The first criterion was to maximize the average performance across patients. This was done by finding the parameter combination (FOI-TOI) which maximized the patient-average AUC for each classifier. The second criterion was to maximize the worst performance across patients. Here, we found the parameter combination (FOI-TOI) which maximized the minimum AUC across patients for each classifier.

### 2.7 Classifier implementation

To further explore the SOZ discrimination potential of nMA and nMS depending on the parameters of interest (FOI-TOI), two distinct methodologies were employed: inter– and intra-patient cross-validation. The aim of both approaches was to establish a patient-specific contact profile and subsequently identify an optimal threshold for distinguishing between SOZ and non-SOZ contacts.

The criterion used to determine this threshold was the F2-score, a variant of the F-beta score with the beta parameter set to 2. The F-beta score, ranging between 0 and 1, serves as a suitable metric for imbalanced data, as it disregards True Negatives (TN) in its calculation, a category often inflated in imbalanced data classification scenarios. In the F-beta score, the beta parameter controls the weight of precision in the score compared to recall. Setting the beta parameter to 2 implies prioritizing recall over precision in the score, and the rationale behind this choice is due to the fact that the relative consequences of a false negative (FN) outweigh those of a false positive (FP) when considering the sizes of both SOZ and non-SOZ groups with respect to the true positives (TP). In essence, setting the beta parameter to 2, we decrease the likelihood of missing a genuine SOZ contact, but at the expense of increasing the risk of misclassifying a non-SOZ contact (Yuan et al., 2024; Semeux-Bernier et al., 2023):

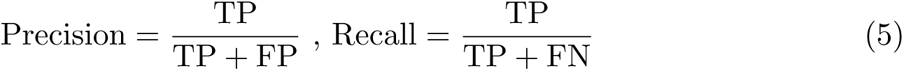

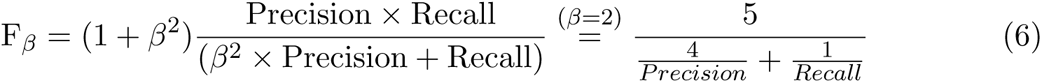

The intra-patient approach was designed to create a specific classification model for each patient, while the inter-patient approach aimed to establish a general model applicable across all patients. The intra– and inter-patient approaches varied in terms of the data utilized for threshold optimization.

In the intra-patient approach, optimization was conducted individually for each patient to seek robust classification across crises: N-1 seizures were utilized as the training set, and their respective nMA and nMS profiles were computed following the methodology described in previous sections. Subsequently, 1000 thresholds were evenly distributed between the minimum and maximum values of all nMA (and nMS) profiles. For each threshold, the F2-score was calculated using the annotated SOZ contacts as the ground truth. The optimal threshold was then determined as the one that maximized the resulting F2-score curve. This optimal threshold was subsequently validated using the F2-score on the tested seizure left out from training. This procedure was iteratively performed for every seizure. The mean testing F2-score value was then calculated, serving as a patient-specific validation metric. Following this, the median across patients was computed to establish the overall cross-validation metric in this approach.

In the inter-patient approach, 9 patients were designated for training, while 1 patient was reserved for testing. The seizure-median nMA and nMS profiles (patient specific) were employed to identify the optimal threshold, following the same F2-score maximizing procedure as described above. Subsequently, this optimal threshold was applied to the test seizure-median profile of the excluded patient. This procedure was iteratively performed, with each patient seizure-median nMA (respectively nMS) profile serving as the test patient while the remaining patients were used for training. The median testing F2-score values across patients was computed to establish the overall cross-validation metric in this approach.

Both methodologies yielded SOZ contact predictions and provided testing validation values across the parameter combinations (FOI-TOI). Therefore, the FOI-TOI combination that maximized the cross-validation testing metric could be selected, along the related optimal threshold for nMA and nMS profiles.

### 2.8 Ethics statement

The study was conducted following the Declaration of Helsinki and informed consent was explicitly obtained from all participants prior to the recordings. All diagnostic and surgical procedures were approved by The Clinical Ethical Committee of Hospital del Mar (Barcelona, Spain).

## 3 Results

### 3.1 Computation of nMA and nMS in the time-frequency space

We analyzed 67 seizures from a total of 10 patients to compare the statistical power and classification performance of a power-based and a connectivity-based biomarker to identify seizure onset contacts across a range of distinct frequencies of interest (FOI) and time windows of interest (TOI). For each seizure and contact, we first computed the time-varying power activation (A) and connectivity strength (S) in each FOI separately. Fig. 2 top shows the time evolution of these metrics in the frequency range 3-160 Hz. We then computed each contact’s normalized mean activation (nMA) and normalized mean strength (nMS) in each TOI. As an example, Fig. 2 bottom shows the nMA and nMS contact profiles in the frequency range 3-160 Hz and over the whole seizure. In general, SOZ contacts consistently displayed increased power (nMA) and decreased connectivity (nMS) values with respect to other contacts for a wide range of FOI-TOI combinations. Increased nMA indicates the presence of enhanced oscillations at the contact, which in turn becomes less synchronized with the rest of the network, as captured by decreased connectivity (nMS). In addition, for this exemplary patient, both nMA and nMS profiles displayed a high degree of similarity across seizures.

**Figure 2.**
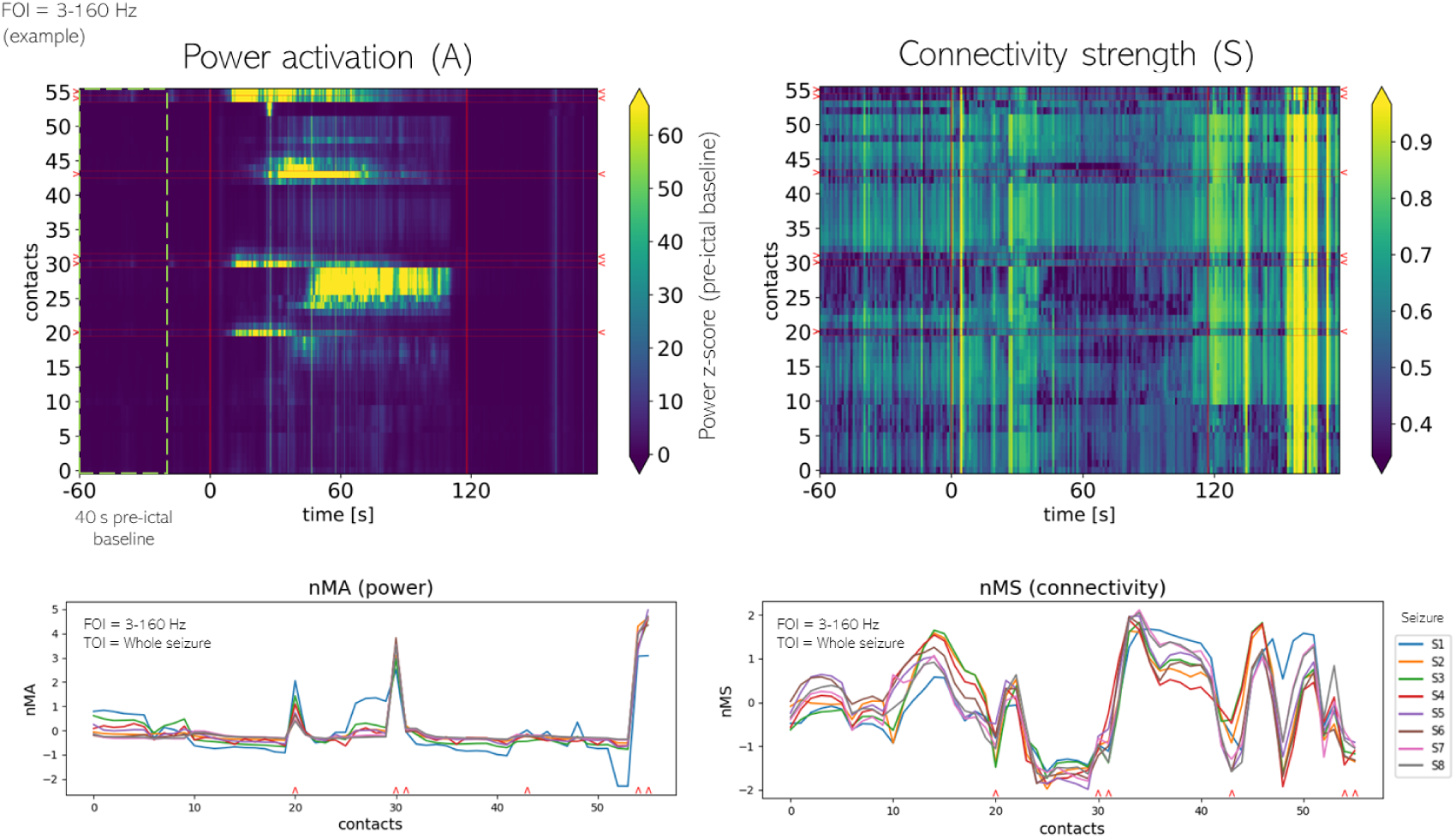
Power– and connectivity-based biomarkers evaluated in 3-160 Hz (FOI) and over the whole seizure period (TOI) in an exemplary seizure of patient P1. Top panels display the broadband (3-160 Hz) time-varying power activation (A, top-left) and connectivity strength (S, top-right). Power values are expressed as a z-score with respect to a baseline distribution built by pooling all contacts’ power values in the time period from –60 s to –20 s. Connectivity strength (S) measures the average functional connectivity of each contact with respect to all other contacts and is computed using a sliding window approach (window length and step of 1 s). Seizure onset and offset times are marked with red vertical lines. The clinically annotated seizure onset zone (SOZ) contacts are marked in red next to the y-axis. Bottom panels display the normalized mean activation (nMA, bottom-left) and normalized mean strength (nMS, bottom-right) computed over the whole seizure period for all seizures of patient P1. Clinically annotated SOZ contacts are marked in red on the x-axis. As previously shown with nMA, the nMS profile displays a high degree of similarity across seizure. SOZ contacts consistently display increased power (nMA) and decreased connectivity (nMS) with respect to other contacts.

### 3.2 Assessing seizure similarity within patients

To leverage the data obtained from several recorded seizures for designing a SOZ biomarker, it is relevant to assess the degree of variability of each variable (nMA, nMS) across seizures in each patient. Here, inter-seizure power (resp. connectivity) similarity was evaluated in each patient and FOI by computing the average Pearson correlation of nMA (resp. nMS) multivariate profiles across all pairs of seizures over the whole seizure period (Fig. S2, top). Inter-seizure similarity was shown to be strongly patient-specific. All patients had large nMA and nMS similarities (*>* 0.6), except for patients P6 and P10. Interestingly, although patient P6 had a medium nMA similarity (0.37 ± 0.06, mean ± SEM), its nMS similarity remained large (0.76 ± 0.03, mean ± SEM). Patient P10 had similar nMA and nMS similarities (0.52±0.03, 0.558±0.018, respectively, mean ± SEM). In addition, nMA and nMS inter-seizure similarity remained high and comparable across frequency bands (≈ 0.8), except in the gamma range of the spectrum, where nMA similarity dropped (0.6 ± 0.1, mean ± SEM, in high-*γ*) while nMS similarity remained stable and high (*>* 0.8). This reinforces the idea that high-frequency nMA may be associated with local neural activity (Vila-Vidal et al., 2023; Buzsáki et al., 2012). Thus, it is more likely to capture the specific electrical patterns of each seizure within patients.

### 3.3 Exploring parameter-dependent SOZ discrimination

After guaranteeing a sufficient degree of seizure similarity, we computed the median nMA (resp. nMS) across seizures and tested for statistical differences between SOZ and non-SOZ contacts using each variable. This was initially illustrated using broadband signals and the whole seizure period (Fig. 3). In this example, nMA (nMS) differences were statistically significant in 8 (resp. 5) out of 10 patients. When significant, differences exhibited very large effect sizes (*D >* 1). In patients P2 and P9, neither nMA nor nMS showed statistically significant differences between SOZ and non-SOZ contacts.

**Figure 3.**
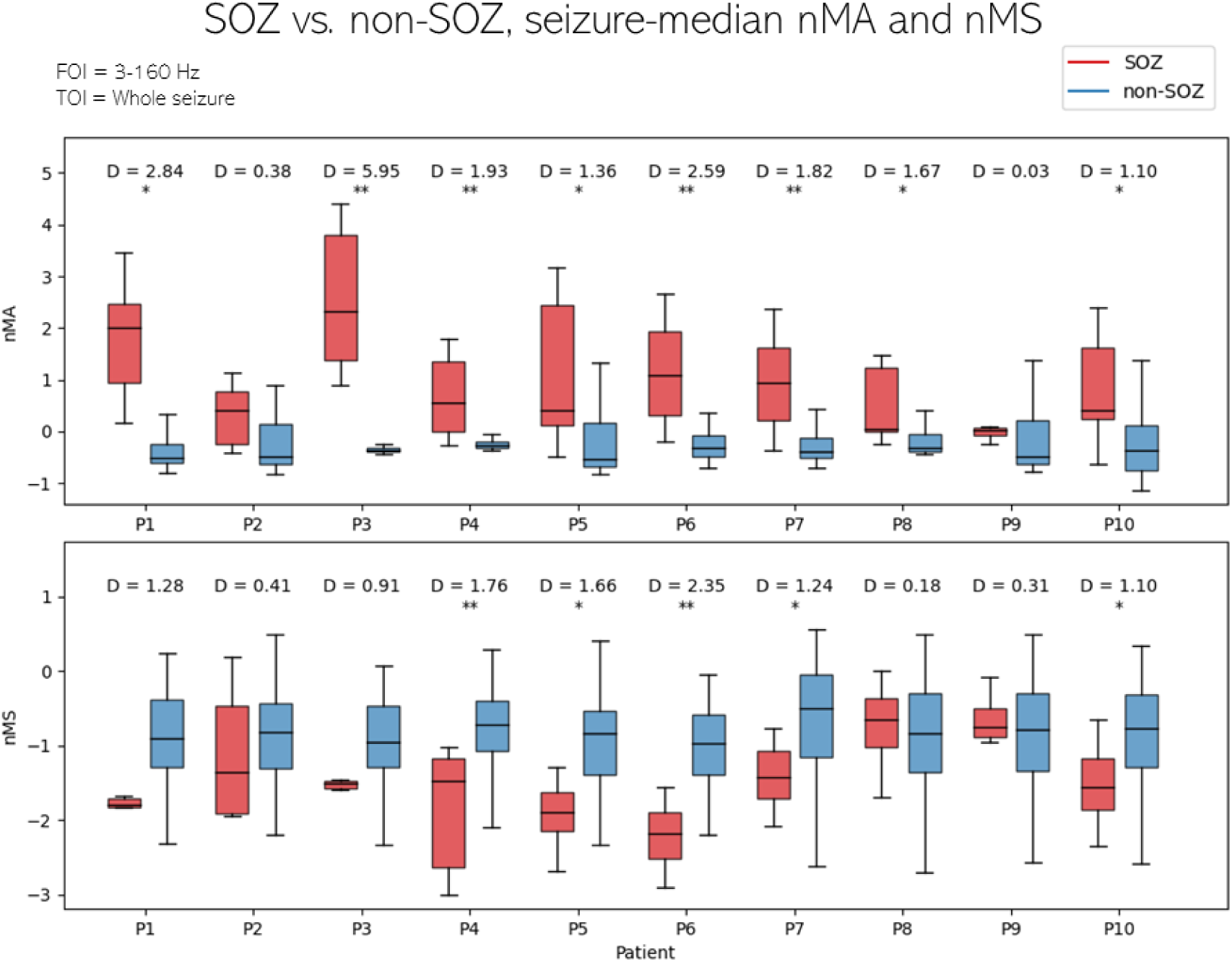
SOZ discrimination with nMA and nMS for an example of parameters of interest (FOI = broadband, 3-160 Hz, TOI = whole seizure period). Boxplots showing the distribution of seizure-median nMA (top) and nMS (bottom) values across SOZ and non-SOZ contacts for each patient. nMA and nMS were estimated for each contact and seizure of patient P1 in 3-160 Hz (FOI) and over the whole seizure period (TOI). Seizure-median nMA and nMS values were obtained for each contact. Differences between SOZ and non-SOZ contacts were tested for statistical significance using a Wilcoxon rank-sum test (** P *<* 0.001, * P *<* 0.01) with multiple-comparison correction using Holm-Bonferroni step-down procedure (Holm, 1979). Effect sizes were evaluated using Cohen’s *d* (reported above each comparison). For this combination of frequency band and time window of interest, all nMA differences were statistically significant (P *<* 0.01) in all cases except in patients P2 and P9. nMS differences were statistically significant (P *<* 0.01) in all cases except in patients P1, P2, P3, P8, P9. When significant, differences exhibited very large effect sizes (*D >* 1). Group (SOZ/non-SOZ) sample sizes for each patient were 6/48, 9/58, 6/50, 11/67, 8/77, 9/90, 11/96, 9/86, 4/120, 13/109.

To explore time-frequency combinations yielding significant SOZ discrimination levels, we computed the effect size for each combination of parameters (FOI and TOI) and extracted the mean and standard deviation group values across patients. As shown in Fig. 4, the nMA (power) exhibited on average larger effect sizes with higher variability than nMS (connectivity) for most FOI-TOI combinations. For nMA (Fig. 4, top), the combinations of *β* band within the range 0-20 s and low-*γ* band up to 0-5 s maximized the patient-average effect size between SOZ and non-SOZ contacts while keeping moderate variability levels. For nMS (Fig. 4, bottom), the combination of high-*γ* band within the range 0-30 s maximized the patient-average effect size between SOZ and non-SOZ contacts but it also yielded the least consistent effect size over the analyzed patients. For completeness, we show in Supplementary Information (Fig. S3) the nMA and nMS effect sizes for each patient.

**Figure 4.**
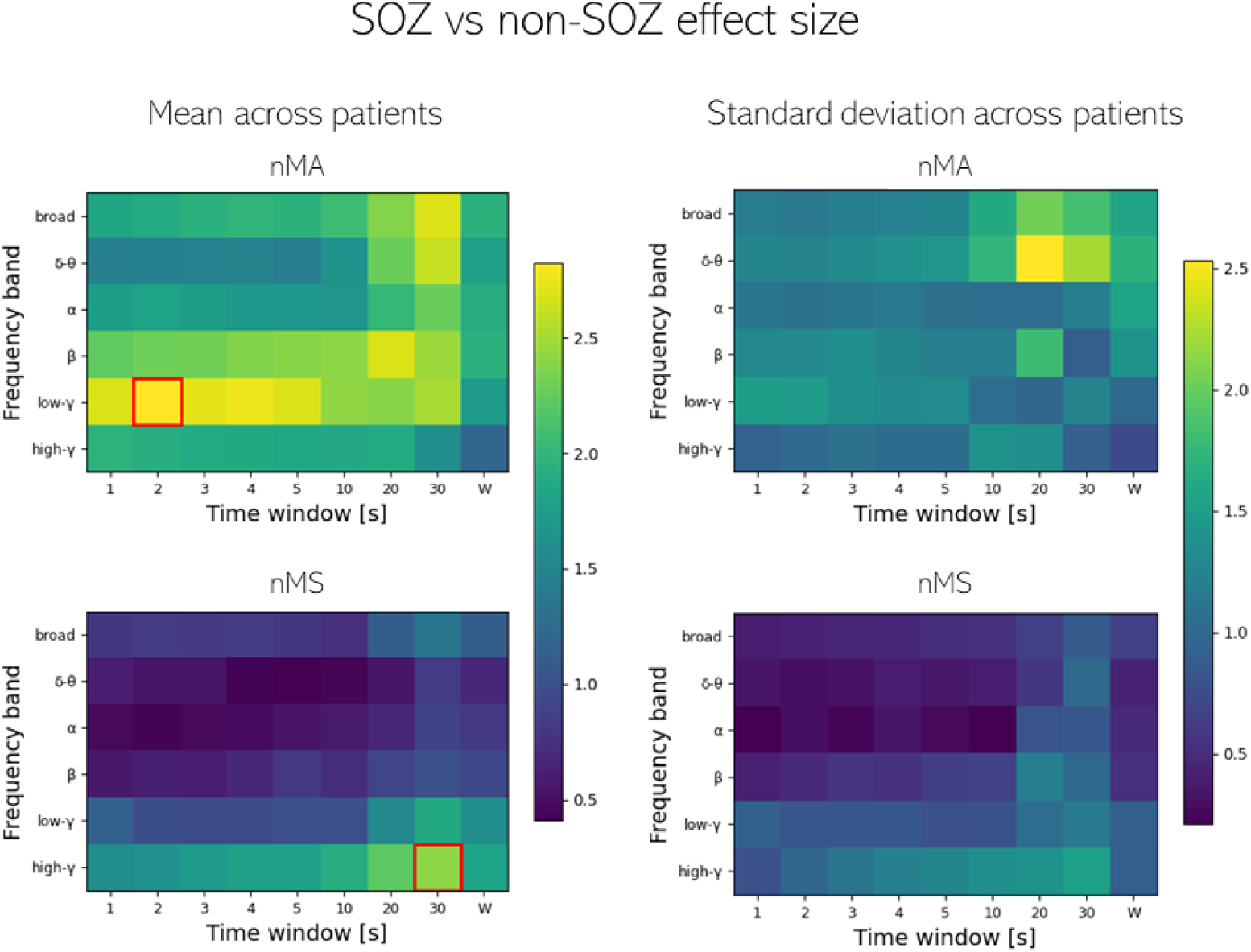
SOZ discrimination (effect size) across patients with nMA and nMS as a function of different FOIs and TOIs. nMA and nMS were estimated for each contact and seizure, and for each combination of FOI (broadband: 3-160 Hz, *δ*-*θ*: 3-8 Hz, *α*: 8-12 Hz, *β*: 12-30 Hz, low-*γ*: 30-70 Hz, and high-*γ*: 70-160 Hz) and TOI (0-1s, 0-2s, 0-3s, 0-4s, 0-5s, 0-10s, 0-20s, 0-30s, and whole seizure period). Seizure-median nMA and nMS values were then obtained for each contact, FOI and TOI. The size of differences between SOZ and non-SOZ contacts with nMA and nMS was quantified using Cohen’s *d* in each FOI and TOI independently. This procedure was repeated for each patient, thus obtaining a distribution of 10 effect size values (one per patient) for each FOI and TOI. Color maps show the mean (left) and standard deviation (right) of nMA (top) and nMS (bottom) effect sizes across patients for every FOI (y-axis) and TOI (x-axis). nMA (power) exhibited on average larger effect sizes with higher variability than nMS (connectivity) for most FOI-TOI combinations. For nMA (top), the combinations of *β* band within the range 0-20 s and low-*γ* band up to 0-5 s maximized the mean effect size between SOZ and non-SOZ contacts. For nMS, the combination of high-*γ* band within the range 0-30 s maximized the mean effect size between SOZ and non-SOZ contacts. See Supplementary Information (A) for patient-specific nMA and nMS effect sizes.

Overall, in Fig. 4 we generalized the patient-level statistical analysis performed in (Vila-Vidal et al., 2017) to a larger frequency-time space and we applied it to each variable (nMA and nMS) independently. This parallel analysis allowed for a fair statistical comparison between nMA and nMS, which might be instrumental to assess the potential of each biomarker to discriminate the SOZ when integrated into classification-based algorithms.

### 3.4 Classification performance between nMA and nMS

Based on the outcomes of the statistical inference, we asked ourselves how the reported patient-average effect sizes might be translated to proper levels of SOZ discrimination when using practical approaches relying on binary classification algorithms.

To address the above question, we quantified and compared the accuracy of a binary classifier based on the nMA and nMS variables, respectively. In brief, for each variable, patient and combination of parameters (TOI-FOI), we computed the area under the curve (AUC) to measure the classifier’s performance. Optimization across parameters’ values was achieved under two different criteria. The first criterion relied on maximizing the patient-average performance. Hence, the patient-average AUC was computed for each classifier over all TOI-FOI combinations (Fig. S4A, left column). In this case, the maximum (squared in red) was attained in the *β* band over 0-30 s for the nMA classifier (0.94 ± 0.02, mean ± SEM across patients) and in low-*γ* band over 0-30 s for the nMS classifier (0.86 ± 0.04, mean ± SEM across patients).

To guarantee a minimum classification performance satisfied by all patients simultaneously, we defined the second criterion as the maximization of the patient’s worst-case performance. Hence, we represented the minimum AUC across patients for each classifier over all TOI-FOI combinations (Fig. S4A, right column). Using this second criterion, the optimal value (maximizing the patient-minimum AUC) remained at the same TOI-FOI combinations as in the first patient-average criterion: *β* band over 0-30 s for the nMA classifier (0.8, minimum across patients) and low-*γ* band over 0-30 s for nMS classifier (0.6, minimum across patients). The distribution of AUC values across patients obtained from optimal selections in both criteria (Fig. S4B) was not statistically significant between measures nMA and nMS (P=0.26, Wilcoxon rank-sum test).

### 3.5 Classifier implementation

To further explore the classification performance of the studied measures, parameter optimization was performed across a predefined time-frequency grid, utilizing two criteria: inter-patient and intra-patient cross-validation, as explained in subsection 2.7. In interpatient optimization, the objective was to determine the seizure-median nMA (resp. nMS) threshold that maximized the F2-score within a training subset of 9 patients. Subsequently, this threshold’s performance was validated on the remaining patient using the same metric. Conversely, intra-patient cross-validation aimed to identify the nMA (resp. nMS) threshold that maximized the F2-score for *N* –1 seizures within each patient, where *N* is the total number of analyzed seizures for a single patient. This optimized threshold was then tested in the remaining seizure, yielding an average F2-score across seizures per each patient.

The results of the inter– and intra-patient cross-validation are depicted in Figure 5. For the intra-patient approach (Fig. 5A left), an optimal parameter combination (FOI-TOI) was identified as the low-*γ* band over 0-10 seconds for nMA (0.77 ± 0.13, median ± MAD across patients) and the high-*γ* band over 0-30 seconds for nMS (0.72±0.11, median ± MAD across patients). Conversely, the inter-patient approach (Fig. 5A right) yielded an optimal combination of the *β* band over 0-30 seconds for nMA (0.82 ± 0.08, median ± MAD across patients) and the high-*γ* band over 0-30 seconds for nMS (0.72±0.12, median ± MAD across patients). In Figure 5B, sensitivity and specificity values are illustrated for the optimal parameter combination obtained from the inter-patient approach. For nMA, sensitivity and specificity were 0.85 ± 0.04 and 0.91 ± 0.03, mean ± SEM across patients, respectively; while for nMS, sensitivity and specificity were 0.65 ± 0.11 and 0.9±0.03, mean ± SEM across patients. Specificity consistently showed high levels across all patients, but sensitivity remained weak, particularly in challenging cases. However, the distribution across patients resulting from optimal selections in both criteria (Figure 5B) showed no statistically significant difference between nMA and nMS in terms of sensitivity and specificity (0.17 and 0.71 respectively, Wilcoxon rank-sum test p-values). For completeness, we show the classification performance of nMA and nMS at the patient level for the intra-patient cross-validation approach in Fig. S5. The best parameter combination for nMA was found to be patient-specific. In contrast, the best frequency band for nMS remained slightly stable across patients in the low– and high-*γ* range. This, together with the increased inter-seizure nMS similarity (Fig. S2), indicates that the results might generalize better for nMS than nMA for larger datasets. The best time window size was also found to be patient-specific, although the majority of the identified optimal time windows extended beyond the range of 0-5 seconds.

**Figure 5.**
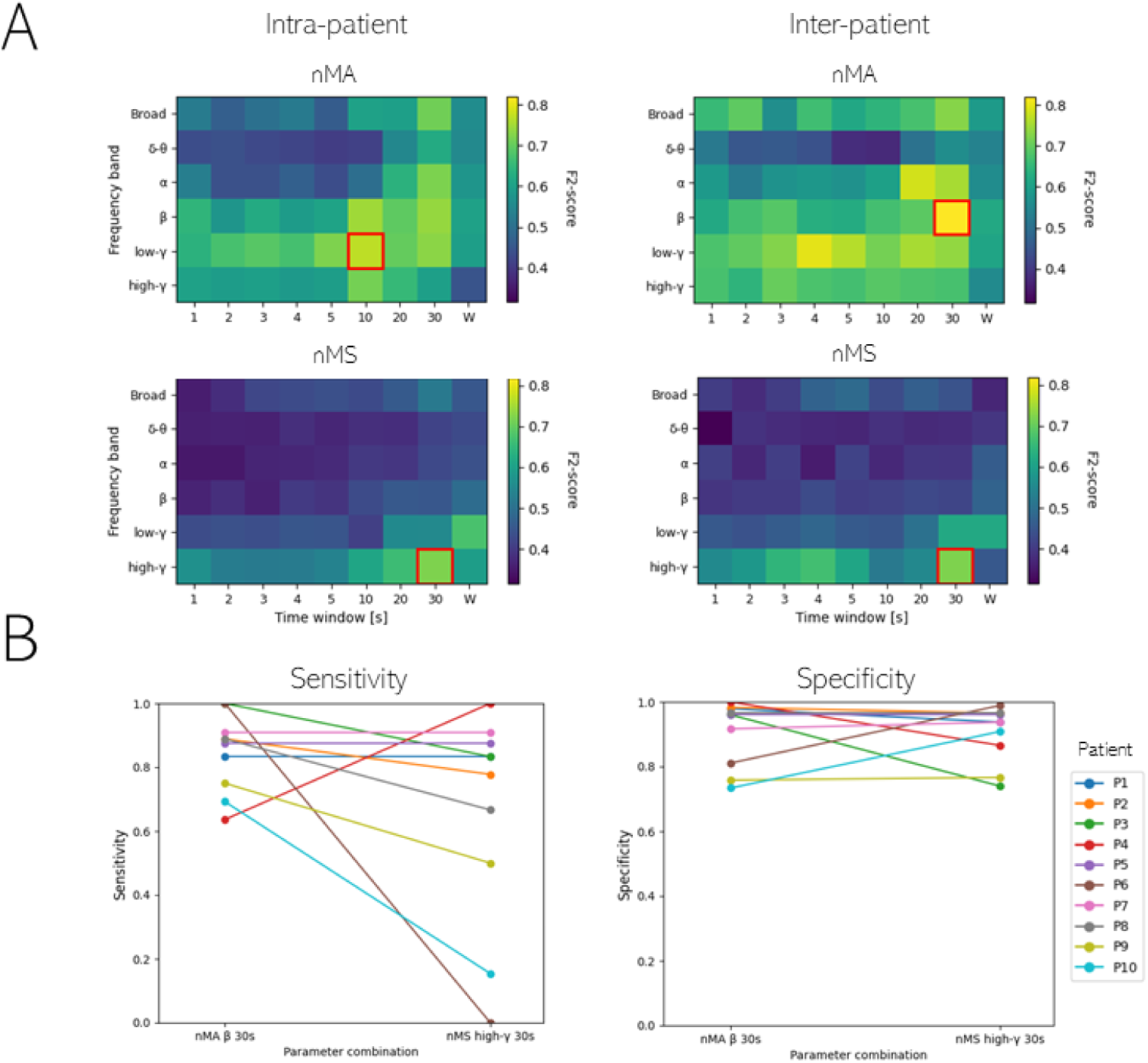
Inter– and intra-patient cross-validation of a SOZ classifier for different FOIs and TOIs. **A**. Color maps illustrate the intra-patient (left) and inter-patient (right) median F2-score values across all validated patients using nMA (top) or nMS (bottom). F2-score ranges between 0 and 1. The optimal test F2-score values for nMA were identified in the low-*γ* band spanning 0-10 s through intra-patient cross-validation, while for inter-patient cross-validation, they were found in the *β* band covering 0-30 s. In the case of nMS, both approaches yielded the same optimal parameter combination: the high-*γ* band over 0-30 s. **B.** Sensitivity (left) and specificity (right) values obtained with the inter-patient maximizing FOI-TOI combination: *β* band and 0-30s for nMA, and high-*γ* band and 0-30s for nMS. Sensitivity and specificity differences across nMA and nMS classifiers were not statistically significant (P *>* 0.05, Wilcoxon rank-sum test).

In general, the results of the classification accuracy under each criteria (AUC patient-average and minimum and F2-score intra– and inter-patient cross-validation criteria) showed that the nMA outperformed nMS for most of the TOI-FOI combinations. However, when selecting the optimal combination for each variable, we did not find significant differences in AUC or sensitivity and specificity between nMA and nMS in our dataset (n=10, P*>*0.1) regardless of the employed criterion.

In conclusion, the application of the proposed methodology to our available dataset suggests that power-based variables might yield slightly better SOZ identification performance than (single-site) functional connectivity variables such as the connectivity strength. Yet, larger datasets will be needed to characterize such comparison more accurately and to investigate whether the combination of each type of variable can also yield performance improvements. In any event, our methodology may be generalized to provide an in-depth and fair comparison analysis between any SOZ discrimination biomarker candidate defined at a single-site level with respect to standard power-based variables.

### 3.6 Application of the algorithm in a practical example

After evaluating and cross-validating the algorithm’s performance using various criteria, we present a case example illustrating how the algorithm could automatically detect the SOZ in a new patient by optimizing FOIs, TOIs, and thresholds over a dataset formed by previous documented patients. In Figure 6A, annotated seizure recordings from a dataset comprising 9 patients are utilized for optimization. Consequently, their respective seizure-median nMA and nMS profiles are extracted, and the optimal parameter combination, along with the associated threshold, is determined using the inter-patient approach using annotated SOZ. In the example, nMA in the training dataset yielded an optimal parameter combination of *β* frequency band, a 0-20 s time window, and the associated nMA threshold of –0.018, while in the case of nMS the optimal parameter combination resulted in high-*γ* frequency band, a 0-30 s time window, and the associated nMS threshold of –1.668.

**Figure 6.**
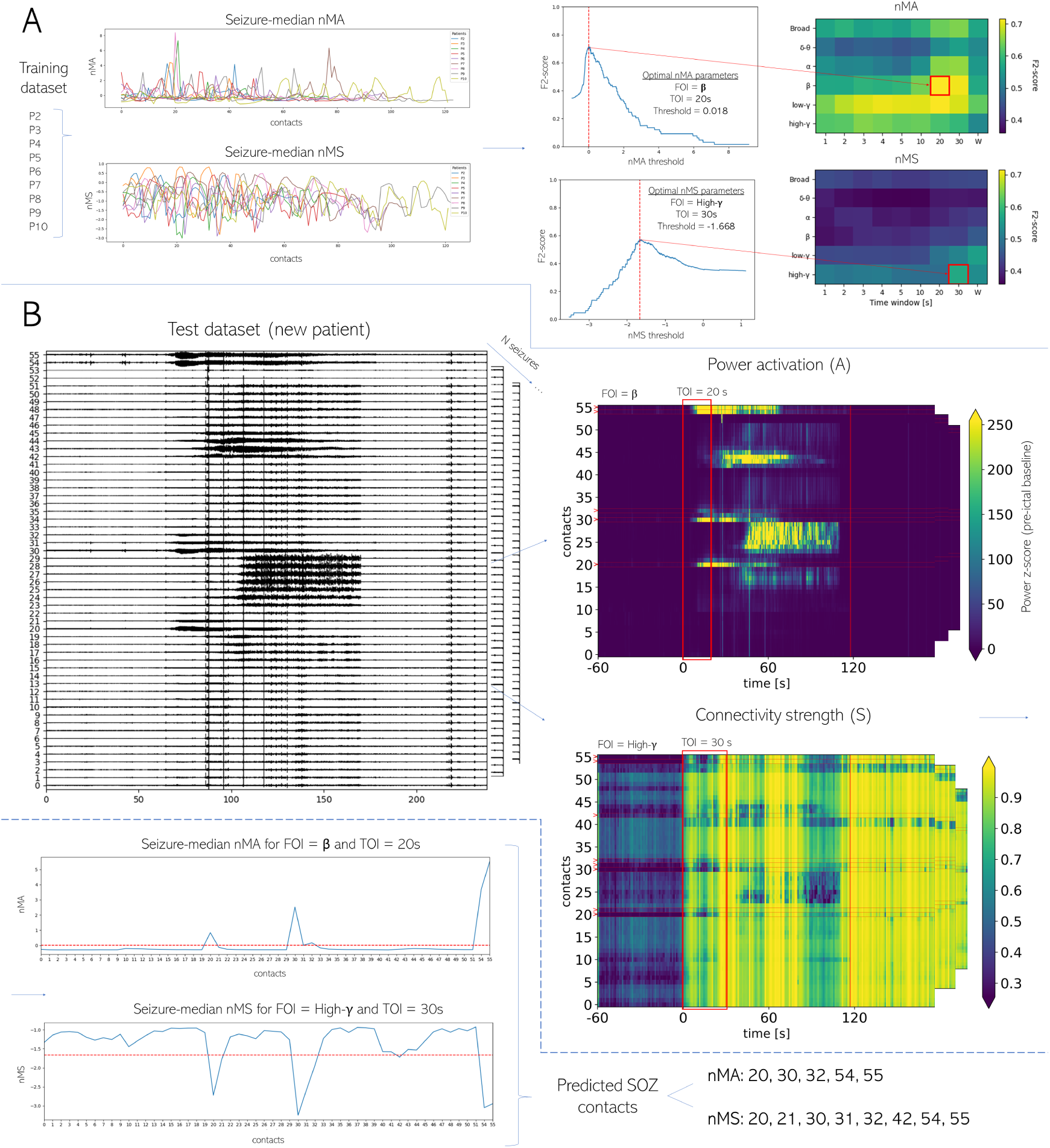
Example case of the inter-patient optimization algorithm. **A**. Median seizure nMA (respectively, nMS) profiles are computed from the training dataset comprising seizures from all training patients. Subsequently, an optimal nMA (respectively, nMS) thresh-old, frequency band of interest (FOI), and time window of interest (TOI) are determined by maximizing the F2-score using annotations of the seizure onset zone (SOZ) within this training dataset. This optimized combination of nMA (respectively, nMS) parameters is then applied to incoming patients. **B.** Upon the arrival of a new patient with N seizures, power activation and connectivity strength plots within the optimal FOI and TOI can be extracted as described in the methodology, and the seizure-median nMA (respectively, nMS) can be computed, using the optimal threshold to identify SOZ contacts. In the illustrated example, patient 1 was used as an unseen testing patient, while all remaining patients composed the training dataset. The sensitivity and specificity achieved for the predicted SOZ contacts, in comparison to the physicians’ SOZ annotations (contacts 20, 30, 31, 43, 54, 55), were 0.67 and 0.98 for nMA, and 0.83 and 0.94 for nMS, respectively.

Subsequently, this optimized parameter combination was applied to the new patient under analysis, depicted in Fig. 6B. Initially, time-varying power activation and connectivity strength maps are visualized using the corresponding optimal frequency band and time window obtained in the previous step. Next, the seizure-median nMA and nMS profiles could be computed, followed by thresholding to identify the SOZ contacts predicted by nMS and nMA. For this specific tested patient, the predicted SOZ contacts using nMA were 20, 30, 32, 54, and 55 (with a sensitivity and specificity of 0.67 and 0.98, respectively, compared to the annotated SOZ), and the predicted contacts using nMA were 20, 21, 30, 31, 32, 42, 54, 55. Sensitivity and specificity values of these predictions when compared to the annotated SOZ (contacts 20, 30, 31, 43, 54, 55) were 0.67 and 0.98 for nMA, and 0.83 and 0.94 for nMS, respectively.

## 4 Discussion

In this work we have presented a parameter-dependent comparison between a power and a connectivity biomarkers based on intracranial EEG extracted from peri-ictal periods for SOZ identification in patients with pharmacoresistant epilepsy. Motivated by previous works (Kramer et al., 2008; Geier et al., 2015; Ponten et al., 2007), one of the central goals of the proposed analysis method was to investigate whether standard power-based biomarkers (Vila-Vidal et al., 2017, 2020b; Gnatkovsky et al., 2011, 2014; David et al., 2011; Bartolomei et al., 2008a) can be outperformed by measures relying on second-order signals’ statistics such as measures derived from delayed functional connectivity.

To study the power-connectivity comparison, we here resorted to a rather general measure, the normalized mean activation, (nMA) (Vila-Vidal et al., 2017) as a proxy of a power-based biomarker, and defined an analogous measure, the normalized mean strength (nMS), as a proxy of a connectivity-based biomarker. The nMS corresponds to the mean functional connectivity (cross-correlation) of every contact’s signal with the rest of signals over a given time window and frequency band, and it can be therefore compared straightforwardly with the nMA. Importantly, the method may be analogously applied to other connectivity or topological measures (Friston, 2011; Bullmore and Sporns, 2009) provided that they are defined at a single-site level (i.e., they can be regarded as vectors of a *n*-space where *n* is the number of contacts). For each selected measure, the method validates that normalized measures are consistent across seizures within a single patient and extracts a meaningful median value (over seizures) that is then used to quantify the effect sizes and classification performance of the SOZ contacts over a grid of relevant physiological frequency bands and nested time windows following the seizure onset.

We illustrated the application of our method to analyze the power-connectivity comparison in a dataset of 10 patients with pharmacoresistant temporal lobe epilepsy that accounted for a total of 67 seizures and whose SOZ was clinically validated by epileptologists. The outcomes of our analysis suggest that power-based biomarkers generally perform better than connectivity-based ones. Yet, our results also show that both biomarkers may achieve similar performance levels when appropriately optimized over frequency bands and time windows of interest.

### 4.1 Seizure homogeneity as initial step for SOZ discrimination

A critical aspect of our method is that it leverages the data from each patient’s recurrent seizures for a single SOZ identification, thus accumulating statistical power across seizures to perform inference and classification at a patient level. As demonstrated in a previous work for power activation (Vila-Vidal et al., 2017), we here showed that the values of connectivity strength also attained a very low variance across seizures when they were normalized across contacts (Fig. S1). In particular, there were only a few seizures that substantially deviated from the median value of nMA and nMS, respectively, in a few patients. Overall, the degree of similarity between seizures was very high and this trend was shown to be consistent across patients regardless of the employed measure (Fig. S2).

Assessing seizure homogeneity was key in our method to extract a median value per contact across seizures, which could provide a sufficiently good representation of the patient’s ictal activity for each measure. This step was therefore instrumental in performing statistical analysis comparing the activity (power/connectivity) between SOZ and non SOZ contacts per patient (Fig. 3). However, it is due stating that the reported seizure homogeneity of our dataset was possibly favored by the fact that all patients had temporal lobe epilepsy with a clear diagnosed focus. It remained out of the scope of this study to investigate to which degree seizure homogeneity still holds in patients with extra-temporal lobe epilepsy or patients with heterogeneous seizures exhibiting more than one focus across different brain areas.

### 4.2 High activation and low functional connectivity identify SOZ

By inspecting the distribution of nMA and nMS values over patients, we generally reproduced known results in the literature (Fig. 3) pointing to the fact that the SOZ is characterized by significantly higher power activation (Vila-Vidal et al., 2017; Bartolomei et al., 2008a) and lower functional connectivity following seizure onset (Kramer et al., 2008). However, this characterization was not uniform among patients when using rather general conditions such as analyzing broadband signals over the whole seizure period. This motivated us to explore how the discrimination of SOZ varied as a function of clinically relevant frequency and time windows. Indeed, the comparison between nMA and nMS over a grained space of parameters allowed us to investigate in depth its dependence on frequency and time. In particular, we observed that for most of the patients, the SOZ discrimination of nMA was mostly maximized in the low-gamma regime (30-50Hz) over the initial 5 seconds (Fig. 4), which reproduced previous own results (Vila-Vidal et al., 2017). With regard to nMS, we observed that the effect size was likely to increase from low to high frequency and from low to higher time window sizes and it achieved its maximum at the high-gamma band within a time window of 30s following seizure onset. Two factors might explain this frequency and time gradient. First, the frequency increase indicates that functional connectivity is a more representative measure of neural activity when inferred at high-frequency bands since it is known to capture interactions across local neural population from different brain regions (Vila-Vidal et al., 2023; Buzsáki et al., 2012). On the other hand, the temporal increase might reflect that functional connectivity variables operate at a smaller timescale than power-based variables over ictal epochs and they thus require larger time window for estimation.

### 4.3 Power-based vs. connectivity-based biomarkers for SOZ classification

Upon reviewing the outcomes of our analysis, we may pose again the question of how connectivity-based biomarkers for SOZ identification compare to power-based biomarkers. Specifically, our results based on the (low computational cost) measure of crosscorrelation reinforce the idea that connectivity measures do not suffice to outperform traditional biomarkers relying on spectral features. Our interpretation behind this behavior is that delayed statistical dependencies, at least at a linear level, do not necessarily reflect information of a pure physiological nature such as propagation pathways, but rather they are a measure of signal similarity. This is specially manifested in the SOZ whose connectivity is lower than in the rest of contacts in the gamma band when the ictal spectral patterns become more differentiated from the rest of the signals after seizure onset. Furthermore, detecting seizure propagation across contacts is not a priori an easy task given the limited spatial sampling of sEEG implantation. Hence, further analysis relying on estimations over observed but also non-observed (latent) data are needed to elucidate how to design connectivity-based biomarkers that can capture patterns associated with seizure propagation, thus complementing the information provided by spectral analysis.

### 4.4 Parameter optimization strategies for SOZ classifiers

This study employed four distinct criteria to select the optimal frequency band and time window for optimizing the SOZ identification performance of the power– and the connectivity-based biomarkers. The initial two methods focused on maximizing the patient-average and patient-minimum AUC, bypassing the need for setting a discrimination threshold. Conversely, the remaining two measures aimed to determine an optimal threshold by maximizing the F2-score in training data and subsequently testing this threshold on separate test data. These latter approaches varied in whether the optimization was conducted to generalize the SOZ discrimination over different seizures within a patient (intra-patient) or it was conducted to generalize the across-seizure average threshold per patient to other patients with the same type of epilepsy (inter-patient).

When comparing criteria for each biomarker independently, both AUC approaches resulted in identical optimal parameter combinations: 30-second time window along with *β* band for nMA or low-*γ* band for nMS. This suggests that the frequency bands and time windows maximizing classification performance on average across all patients are also optimal parameter combinations for the patient with the lowest AUC. Conversely, the binary classification of nMA and nMS relying on F2-score showed a slightly different ensemble of optimal time-frequency combinations. In particular, for nMA, the generalization over patients stayed at the beta band and 30 seconds time window, while generalization across seizure per patient shifted to the low-gamma band and 10-seconds time window, despite maintaining good performance in the former combination.

Furthermore, it is noteworthy that, when taking all four criteria into account, the optimal parameter combination for the power-based measure, considering all patients collectively, resulted in the *β* frequency band and a 30-second time window. Conversely, the optimal parameter combination for the connectivity-based SOZ classifier was identified within the *γ* frequency band and the same 30-second time window, regardless of whether patients were assessed as a group or individually. These results may indicate that the strategies employed to identify the optimal frequency and time settings for all patients are comparable. Consequently, they could serve as a reliable approach for optimizing classification performance tasks across various parameter combinations.

Moreover, a significant distinction between optimal combinations obtained from cross-validated classification and statistical comparison is the utilization of larger windows. Small windows for nMA proved ineffective in cross-validation, suggesting that achieving robustness in this manner requires the integration of information over an extended recording period.

### 4.5 Considerations for classifier generalization

Considerations must be taken into account when attempting to optimize a general classification model for all patients within an epilepsy type. As demonstrated in this study, the optimal parameter combination may vary among patients (see Fig. S5). Consequently, selecting a single optimal frequency band and time window combination for all patients may yield suboptimal results. The variability across individual optimal combinations can be attributed to several causes, one being the differences between their seizure nMA and nMS contact profiles. In our case, patients 6, 9 and 10 offered poor inter-seizure similarity in their nMA/nMS profiles (Fig. S2 top), which explains the suboptimal results achieved by the general classifier at classifying SOZ on these patients (Fig. 5B and Fig. S4B). Variability might be also explained by different inherent frequency seizure-onset patterns between patients (Vila-Vidal et al., 2020a). In our study this is reflected in the frequency bands that maximize effect sizes for every patient (see Fig. S3 top). To sum up, any source of variability must be carefully regarded when selecting the data that will constitute the training dataset.

However, once the aforementioned variability has been tackled, a potential perspective on generalizing the classification model across patients could be to identify common trends present in temporal epilepsy, distinct from other types, in terms of changes in contact power or functional connectivity. By stratifying group-level patterns, it becomes plausible to delineate characteristic network alterations that are indicative of temporal epilepsy against other types. Identifying commonalities in network alterations not only enhances the understanding of the underlying mechanisms but also holds potential for refining prognostic markers tailored to specific patient cohorts.

### 4.6 Limitations

In this study, we performed a comparison between a power– and a connectivity-based measure to infer the SOZ from sEEG signals. As a result of our analyses, we could establish the optimal parameters for each type of measure in our dataset, but we did not build any joint biomarker that combines both power and connectivity. Further research should investigate how to combine both sources of information, considering that this might vary depending on various factors, such as the epilepsy type, the spectral features of the seizure onset patterns or the spatial extent of the implantation. In addition, it should be noted that our results are based on the assessment of a very specific connectivity measure derived from cross-correlation. Cross-correlation is defined to capture linear relationships between signals with possible time delays. Other connectivity measures might target other association features between signals: nonlinear dependencies, phase-phase couplings or amplitude-amplitude couplings, among others. Some of these measures might capture independent phenomena and, so, our results cannot be generalized.

Concerning the comparison of parameter combinations, such as frequency bands and time windows of interest, there is room for further exploration. Dividing the canonical bands into narrower frequency segments might provide more detailed clinical insights, especially regarding specific bands and their relationship with SOZ. However, a notable limitation arises from the definition of time windows, which were determined starting at the clinically annotated seizure onset. Therefore, further exploration to refine the boundaries of these time windows is necessary.

The major limitation of this work, however, is that the power and connectivity analyses were performed retrospectively after the SOZ was defined by clinical electrophysiological criteria and the compromised contacts were determined ad hoc. To ultimately assess the clinical applicability of these and other measures, in future studies, these localization analyses should be performed prospectively in a blinded fashion with respect to the classically defined SOZ.

## 5 Conclusions

This study demonstrates the importance of optimizing frequency bands and time windows when evaluating power or connectivity based seizure onset zone (SOZ) discrimination biomarkers in ictal stereo-encephalography (sEEG) recordings. By comparing and fine-tuning a power and a connectivity-based biomarkers, we observed that the former typically exhibits superior performance over the latter in identifying seizure onset zone (SOZ) contacts. However, by independently optimizing both biomarkers, maximizing either AUC or F2-score, comparable performance levels can be achieved. Our findings indicate that for the power biomarker, optimal performance is attained in the beta and lower-gamma frequency bands, while for connectivity, the higher-gamma band proves optimal SOZ classification, both within a 30-second window post-seizure onset.

These results underscore the necessity of meticulous optimization over frequency bands and time windows when assessing SOZ discrimination biomarkers. Such insights are crucial for the development of robust SOZ classifiers utilizing retrospective patients’ data for clinical applications. By incorporating this optimization step, clinicians can enhance the accuracy and reliability of SOZ localization, thereby improving patient outcomes in the management of pharmacoresistant temporal lobe epilepsy.

## Acknowledgements

M.V. and R.Z. were funded by the European Regional Development Funds, Grant/Award Number: CECH/001-P-001682. A. T. C. and M.V. were funded by the Spanish Ministry of Science, Innovation, and Universities (MCIU), Grant/Award Number: PID2020-119072RA-I00/ AEI/10.13039/501100011033. G.D. was supported by the AGAUR research support grant (ref. 2021 SGR 00917) funded by the Department of Research and Universities of the Generalitat of Catalunya, the NODYN Project PID2022-136216NB-I00 financed by the MCIN/AEI/10.13039/501100011033/FEDER, UE., the Ministry of Science and Innovation, the State Research Agency and the European Regional Development Fund and the project NEurological MEchanismS of Injury, and Sleep-like cellular dynamics (NEMESIS) (ref. 101071900) funded by the EU ERC Synergy Horizon Europe.

## A Supplementary Information

**Figure S1.**
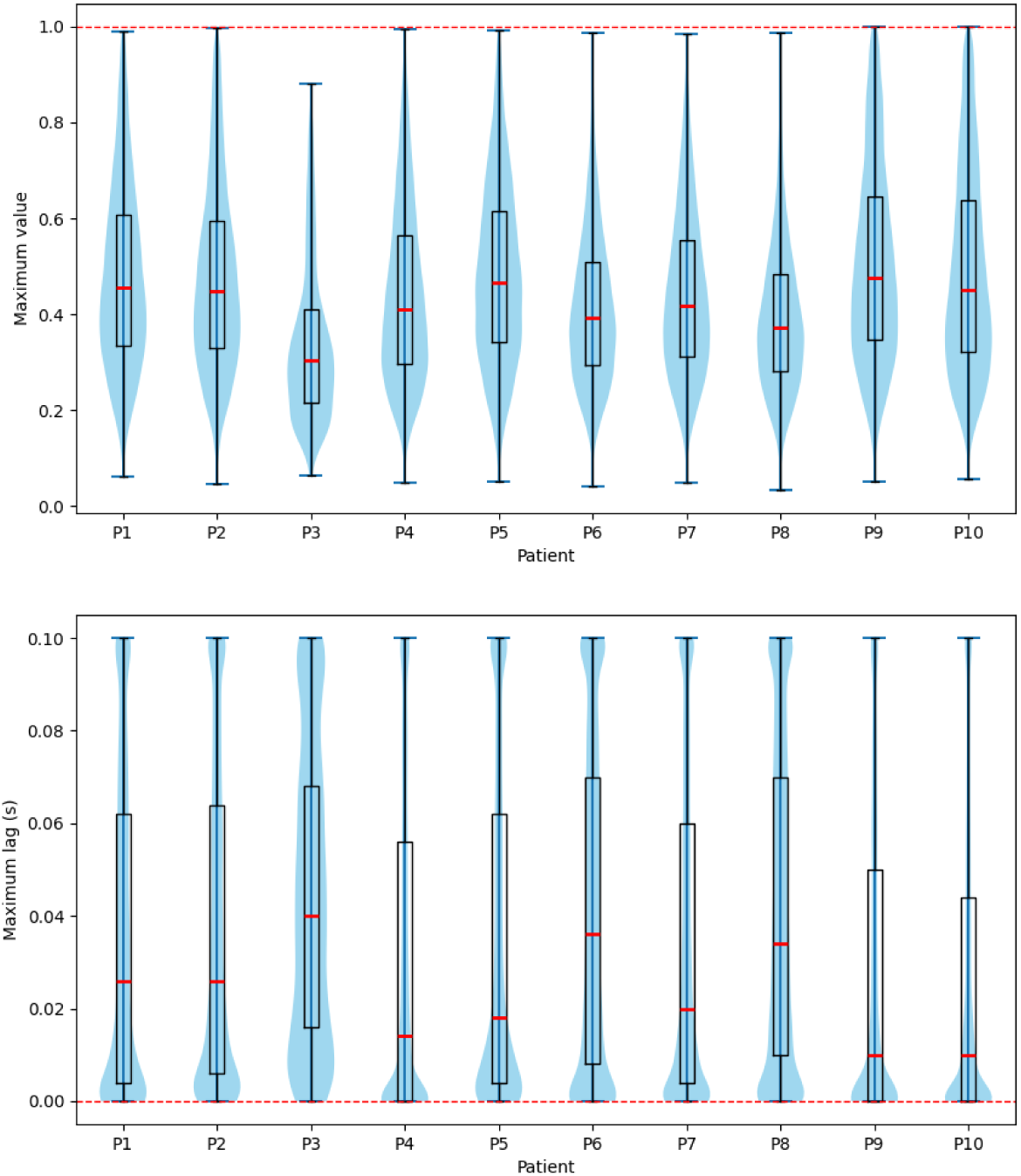
Distribution of pairwise cross-correlation maximum values and lags surrounding seizure onset. The box and violin plots illustrate the patient distributions of SOZ and non-SOZ pairwise cross-correlation maximum values (top) and lags (bottom) within time windows spanning 5 seconds before and after seizure onset. Maximum cross-correlation values are distant from 1, indicating limited correlation between SOZ and non-SOZ contacts. Moreover, although a notable cluster of lags stays near 0, implying immediate maximal correlation among specific contacts, a considerable portion of delays diverges from this point covering the entire spectrum of possible lags. These observations suggest the absence of substantial volume conduction and non-silent reference effects. Otherwise, the majority of maximum values would have been 1, and most lags would have clustered around 0 (red lines). This observation is further evidenced by the exemplary signals and functional connectivity matrix depicted in Figure 1B.

**Figure S2.**
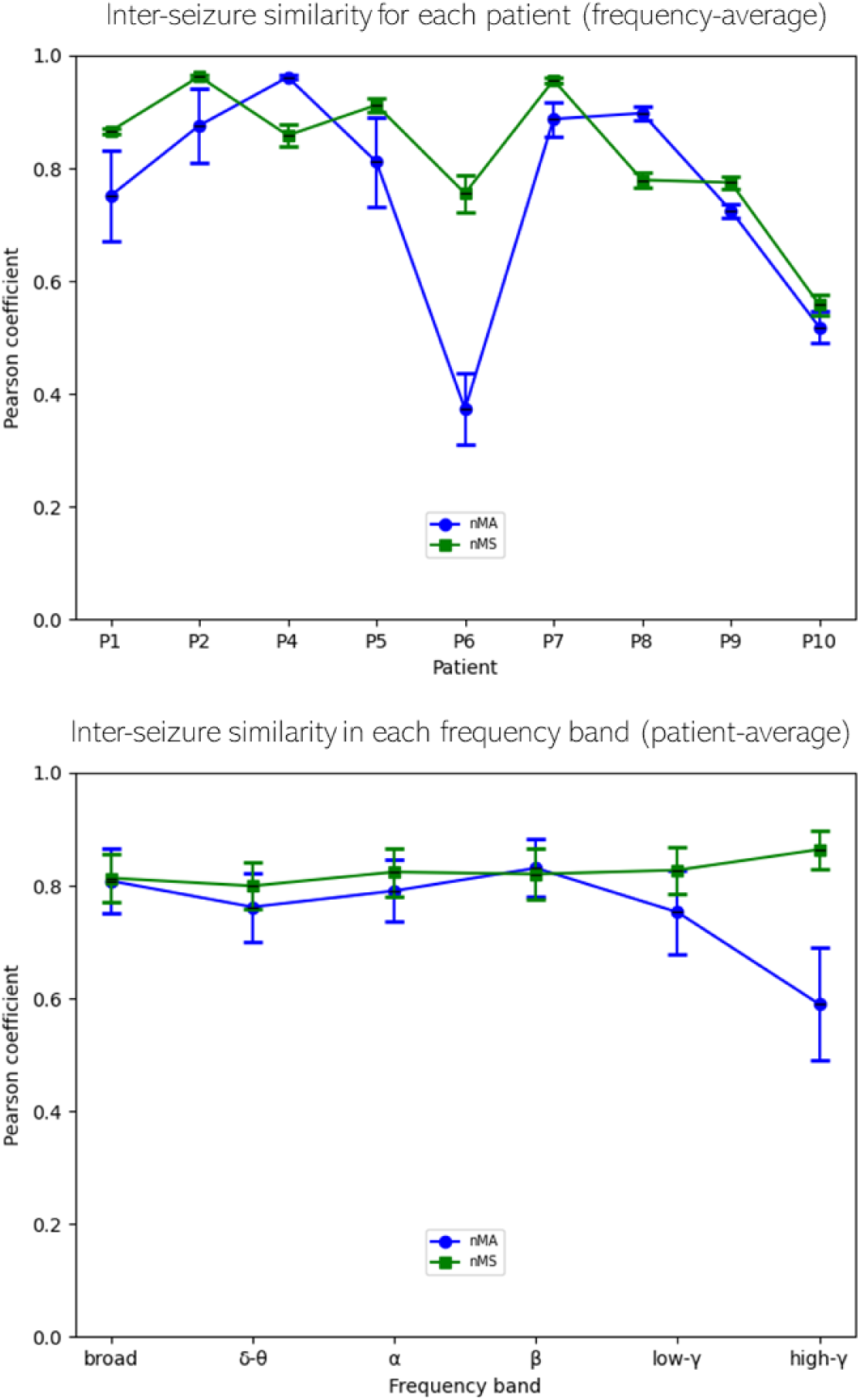
Inter-seizure similarity across patients (frequency-average) and across frequency bands (patient-average). The mean Pearson coefficient was computed to assess the similarity of nMA (resp. nMS) contact profiles across seizures. Similarities across patients with error bars indicating the standard error of the mean related to variability across frequency bands (top) and similarities across frequency bands with error bars related to variability across patients (bottom) are shown for the whole seizure period. Patient 3 was excluded in this figure because it only had one seizure, and therefore no inter-seizure variability could be computed. For inter-seizure variability across patients (top), all patients had mean similarities above 0.5 except for nMA in patient 6. Distribution of similarities across frequency bands (bottom) was similar between nMA and nMS except for high-*γ* frequency bands.

**Figure S3.**
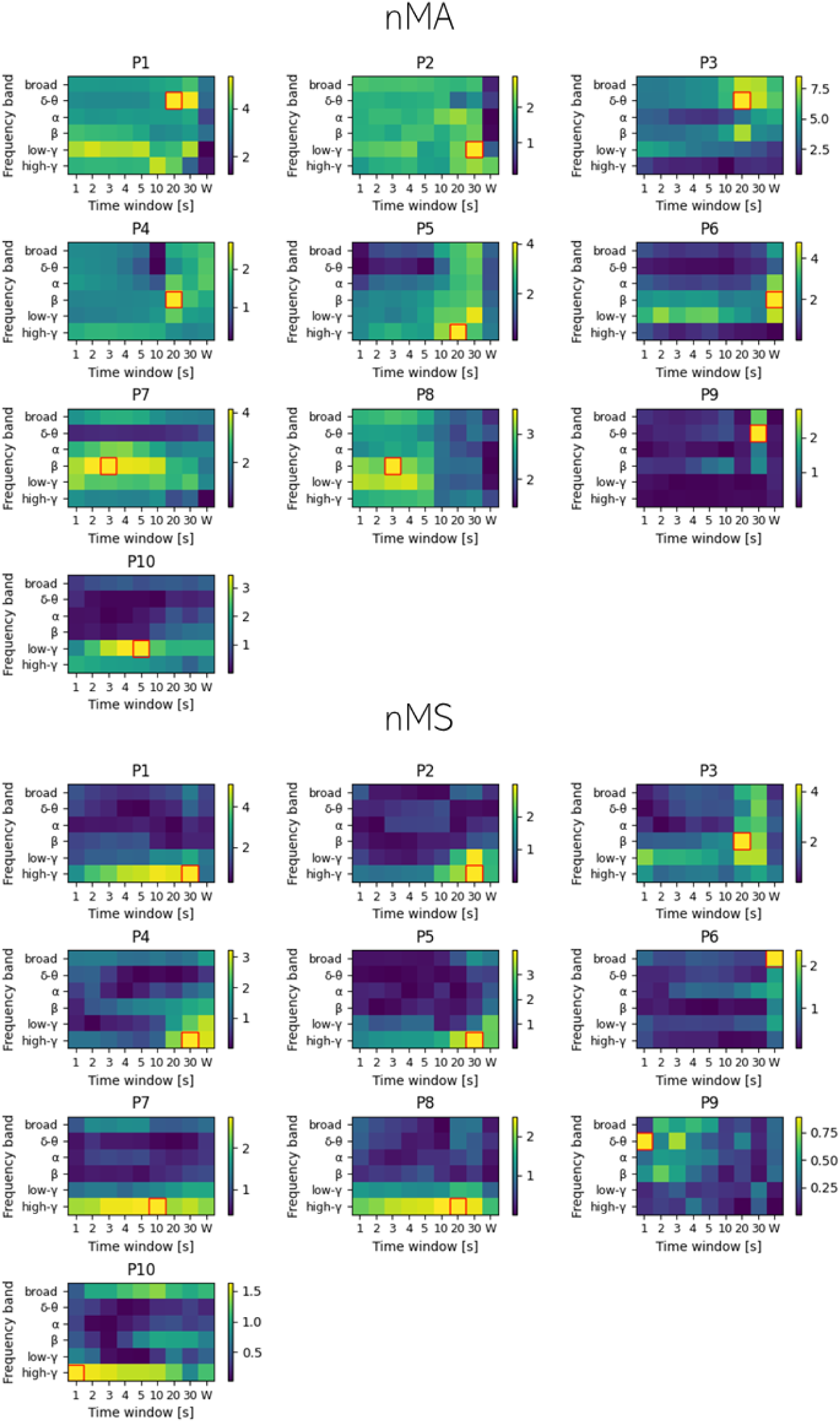
SOZ discrimination (effect size) for each patient with nMA and nMS as a function of different FOIs and TOIs. Color maps show the seizure-median nMS (top) and nMA (bottom) effect size (Cohen’s *d*) values across SOZ and non-SOZ groups for the different frequency bands (y-axis FOIs: broadband: 3-160 Hz, *δ*-*θ*: 3-8 Hz, *α*: 8-12 Hz, *β*: 12-30 Hz, low-*γ*: 30-70 Hz, and high-*γ*: 70-160 Hz) and time windows (x-axis TOIs: 0-1s, 0-2s, 0-3s, 0-4s, 0-5s, 0-10s, 0-20s, 0-30s, and whole seizure pe2ri9od). For nMS (connectivity), high-*γ* frequency band and long time windows (*>* 20 s) maximized effect size in 7 out of 10 patients. Effect size values were greater in nMA (color bar scales) than in nMS for all patients except for patient 4. In nMA, mid-high frequency bands (beta and low-gamma) maximized effect size in 7 out of 10 patients, while effect size-maximizing time windows were different across patients.

**Figure S4.**
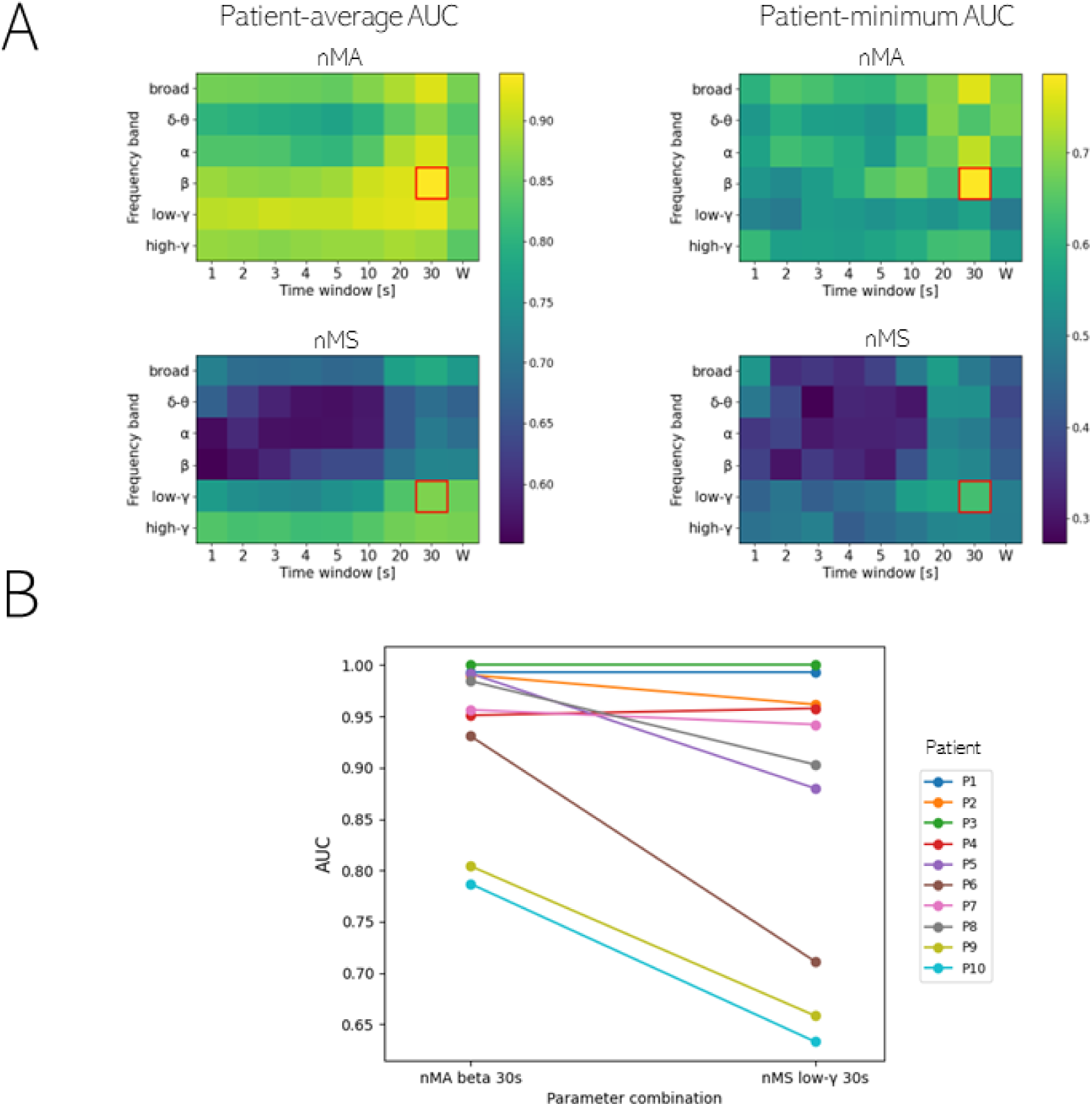
SOZ vs non-SOZ classification performance based on nMA and nMS values. A. Parameter optimization (over a pre-defined time-frequency grid) based on two criteria: patient-average and patient-minimum performance. The optimization criterion used in patient-average performance is based on maximizing the average performance across patients, while the criterion used in patient-minimum performance is based on maximizing the worst-case scenario. Each panel shows patient-average (left) and patient-minimum (right) AUC values for the SOZ classifier based on seizure-median nMA (top) and seizure-median nMS (bottom). The maximum AUC values following each criterion were both found in the *β* band over 0-30 s for nMA, and in low-*γ* band over 0-30 s for nMS. **B.** AUC values obtained in each patient with the nMA and nMS classifiers when using the optimal parameter combinations found in (A). AUC values for the obtained combinations were all above 0.6 for nMS and above 0.75 for nMA. Differences between AUC values were not statistically significant between nMA and nMS (Wilcoxon rank-sum test p-value *>* 0.05).

**Figure S5.**
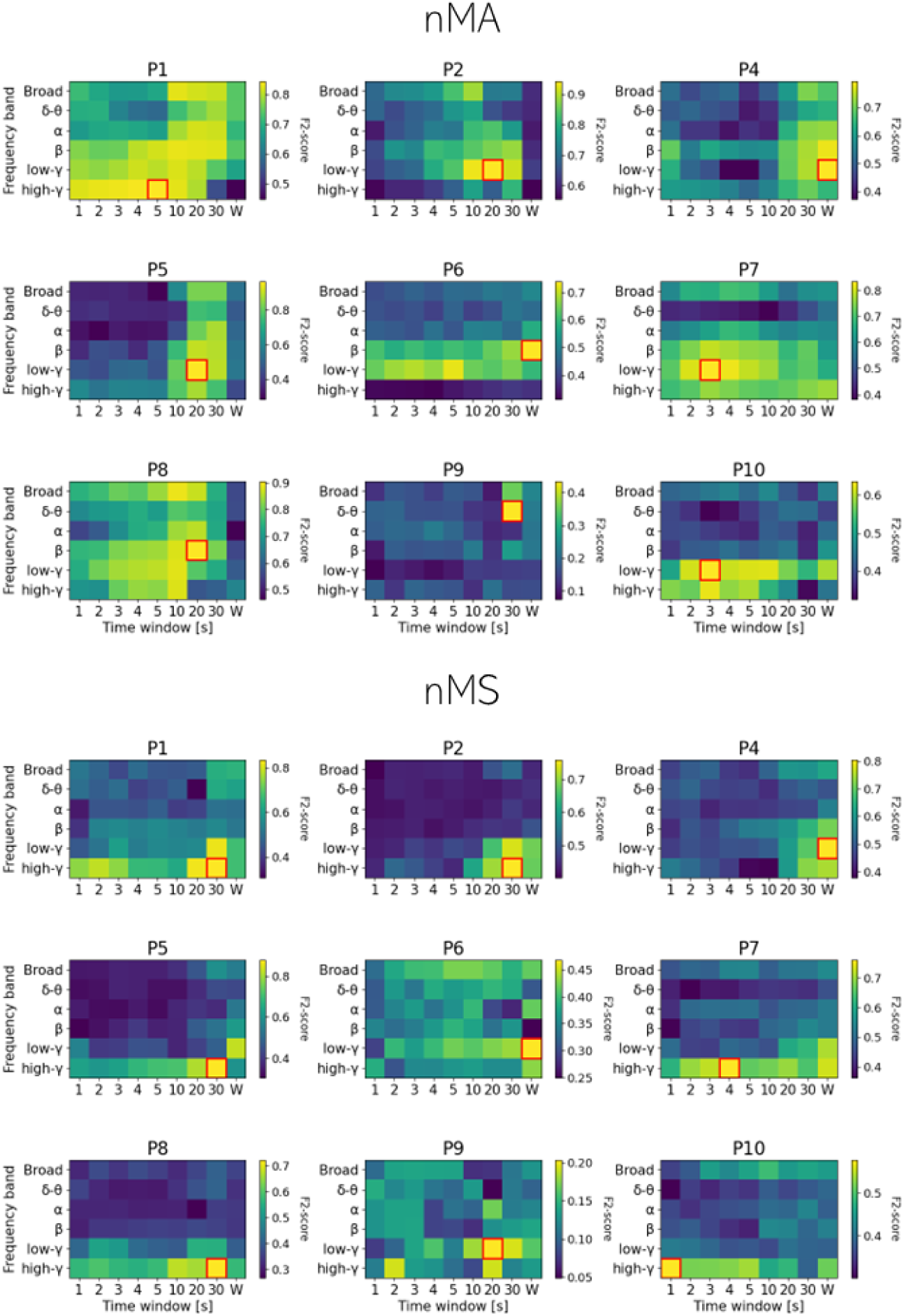
Seizure-Average F2-Score in Intra-Patient Cross-Validation of nMA and nMS. Color maps display the nMA (top) and nMS (bottom) seizure-average F2-score values across SOZ and non-SOZ groups acquired during testing following intra-patient cross-validation. In this procedure, N-1 seizures of each patient were used to optimize an nMA (respectively nMS) threshold by maximizing the F2-score. Subsequently, this threshold was applied to the remaining seizure to obtain a validation F2-score value. Finally, all validation values obtained for each validation seizure were averaged to yield a single validation value for each patient and parameter combination, depicted in the color map grids. Regarding nMA, a frequency of interest (FOI) of high-frequency bands (ranging from *β* to high-*γ* band) was observed to maximize the validation metric in 8 out of 9 patients. Conversely, for nMS, low– and high-*γ* bands along with long time windows of interest (TOI) were identified to optimize the seizure average F2-score in 7 out of 9 patients.

## Notes

### Competing Interest Statement

The authors have declared no competing interest.

### Summary of Updates

We have changed the title of the manuscript to clarify that the presented methodology focuses on two biomarkers and is applied to patients with temporal lobe epilepsy. We have gone beyond the previous AUC analysis to provide a practical implementation of a binary SOZ classifier based on F-score that generalizes results in two different directions: a) over seizures for each individual patient, and 2) over different patients with temporal lobe epilepsy. The classification cross-validation results are now shown in Figure 5 and the implementation is now illustrated in the new Figure 6. We have added a new Figure 1 in Supplementary Information where we prove by means of cross-correlation maximum values and lags that no relevant volume conduction or non-silent reference phenomena were affecting the data, especially in our specific analysis. In addition, in Figure 1 non-neighbour contacts were shown to illustrate the cross-correlation maximum values especially between SOZ and non-SOZ contacts (resulting in low maximum values, i.e., low connectivity, and non-zero lags). We have further justified the choice of the cross-correlation as a good representative of state-of-the-art linear connectivity biomarkers for SOZ discrimination. We have added further information on how the epileptologists annotated the data. We have implemented the Holm-Bonferroni procedure for controlling the family-wise error rates in multiple comparisons, shown in Figure 3. We have implemented a subsection in discussion explaining considerations for the classifier generalization. We have further discussed the study limitations on the definition of the time windows.

